# MAHOMES II: A webserver for predicting if a metal binding site is enzymatic

**DOI:** 10.1101/2023.03.08.531790

**Authors:** Ryan Feehan, Matthew Copeland, Meghan W. Franklin, Joanna S. G. Slusky

## Abstract

Recent advances have enabled high-quality computationally generated structures for proteins with no solved crystal structures. However, protein function data remains largely limited to experimental methods and homology mapping. Since structure determines function, it is natural that methods capable of using computationally generated structures for functional annotations need to be advanced. Our laboratory recently developed a method to distinguish between metalloenzyme and non-enzyme sites. Here we report improvements to this method by upgrading our physicochemical features to alleviate the need for structures with sub-angstrom precision and using machine learning to reduce training data labeling error. Our improved classifier identifies protein bound metal sites as enzymatic or non-enzymatic with 94% precision and 92% recall. We demonstrate that both adjustments increased predictive performance and reliability on sites with sub-angstrom variations. We constructed a set of predicted metalloprotein structures with no solved crystal structures and no detectable homology to our training data. Our model had an accuracy of 90 - 97.5% depending on the quality of the predicted structures included in our test. Finally, we found the physicochemical trends that drove this model’s successful performance were local protein density, second shell ionizable residue burial, and the pocket’s accessibility to the site. We anticipate that our model’s ability to correctly identify catalytic metal sites could enable identification of new enzymatic mechanisms and improve *de novo* metalloenzyme design success rates.

**Significance statement:** Identification of enzyme active sites on proteins with unsolved crystallographic structures can accelerate discovery of novel biochemical reactions, which can impact healthcare, industrial processes, and environmental remediation. Our lab has developed an ML tool for predicting sites on computationally generated protein structures as enzymatic and non-enzymatic. We have made our tool available on a webserver, allowing the scientific community to rapidly search previously unknown protein function space.

## 1. Introduction

Enzymes are biological catalysts that are known to lower activation energy for over 8,000 reactions (McDonald, Boyce, and Tipton 2009). Furthermore, enzymes can increase reaction rates by factors of up to 10^17^-fold (Wolfenden, Ridgway, and Young 1998). Enzymes are also becoming increasingly prevalent in industrial processes due to their greener chemistry (Sheldon and Woodley 2018). Despite the importance and extent of enzymatic research, a reproducible physicochemical basis of catalysis remains elusive. This unknown limits *de novo* enzyme design or even reliable identification of enzyme active sites from structure.

We have recently used protein structure-based machine learning (ML) to distinguish between very similar sites, metalloenzyme active sites and protein sites that bind metals without any enzyme activity (Feehan, Franklin, and Slusky 2021). Our model, <u>m</u>etal <u>a</u>ctivity <u>h</u>euristic <u>o</u>f <u>m</u>etalloproteins and <u>e</u>nzyme <u>s</u>ites (MAHOMES), uses an extra-trees algorithm to achieve better performance metrics than available enzyme function predictors. We attribute the classifier’s success to training on structural physicochemical properties of similar sites. By training on negative sites that were also in pockets and also coordinated metals, rather than on all other sites on the protein, our classifier was able to assign feature importance based on characteristics that were particular to enzyme activity.

Using protein structure-based features enabled MAHOMES to focus on learning physicochemical properties related to catalysis but it relied on structurally determined proteins for its input. The PDB only has ∼200,000 solved protein structures (Burley et al. 2019) thereby limiting MAHOMES utility. Recently, the ML tool AlphaFold2 generated quality protein structure predictions (Jumper et al. 2021) and now two hundred million predicted structures are available for download from AlphaFoldDB (Tunyasuvunakool et al. 2021). However, it remained unclear if these structures could be used for identifying catalytic sites. This concern was compounded by the finding that a relatively low percentage of AlphaFold models have a high enough confidence to be recommended for characterizing binding sites (Thornton, Laskowski, and Borkakoti 2021).

To test usage of the computationally generated structures, we updated the calculation methods used for several of MAHOMES features to reduce the need for sub-angstrom accuracy. Then, we use the new features when cross validating ML models to reduce labeling error in our training data labels. The improved features and training data were used with a variety of ML classifiers and techniques. We found that both the feature improvement and reduced labeling error led to increased performance for ML models. Our best ML model, MAHOMES II, outperformed its predecessor on our holdout test-set with 94% precision and 92% recall. Furthermore, MAHOMES II’s predictions were more reliable for different input structures of the same site with sub-angstrom differences. We evaluated MAHOMES II on a new set of predicted metalloprotein structures, where it scored 97.5% accuracy on high confidence structures. Finally, we examined the features that MAHOMES II found to be the most important for making successful enzyme or non-enzyme predictions and found a preference for features describing enzyme sites to be densely packed, have buried second shell residues, and pockets that were highly accessible to the metal. MAHOMES II can be accessed online (https://mahomes.ku.edu), allowing easy use for the scientific community, regardless of computational expertise.

## 2. Results

### 2.1 New and improved feature calculations

To transform metal sites into input for ML algorithms, we identified features belonging to five categories which have previously been linked to enzymatic activity– coordination geometry, electrostatics, pocket lining, pocket void, and Rosetta energy terms (Figure 1 a and b).

**Figure 1:**
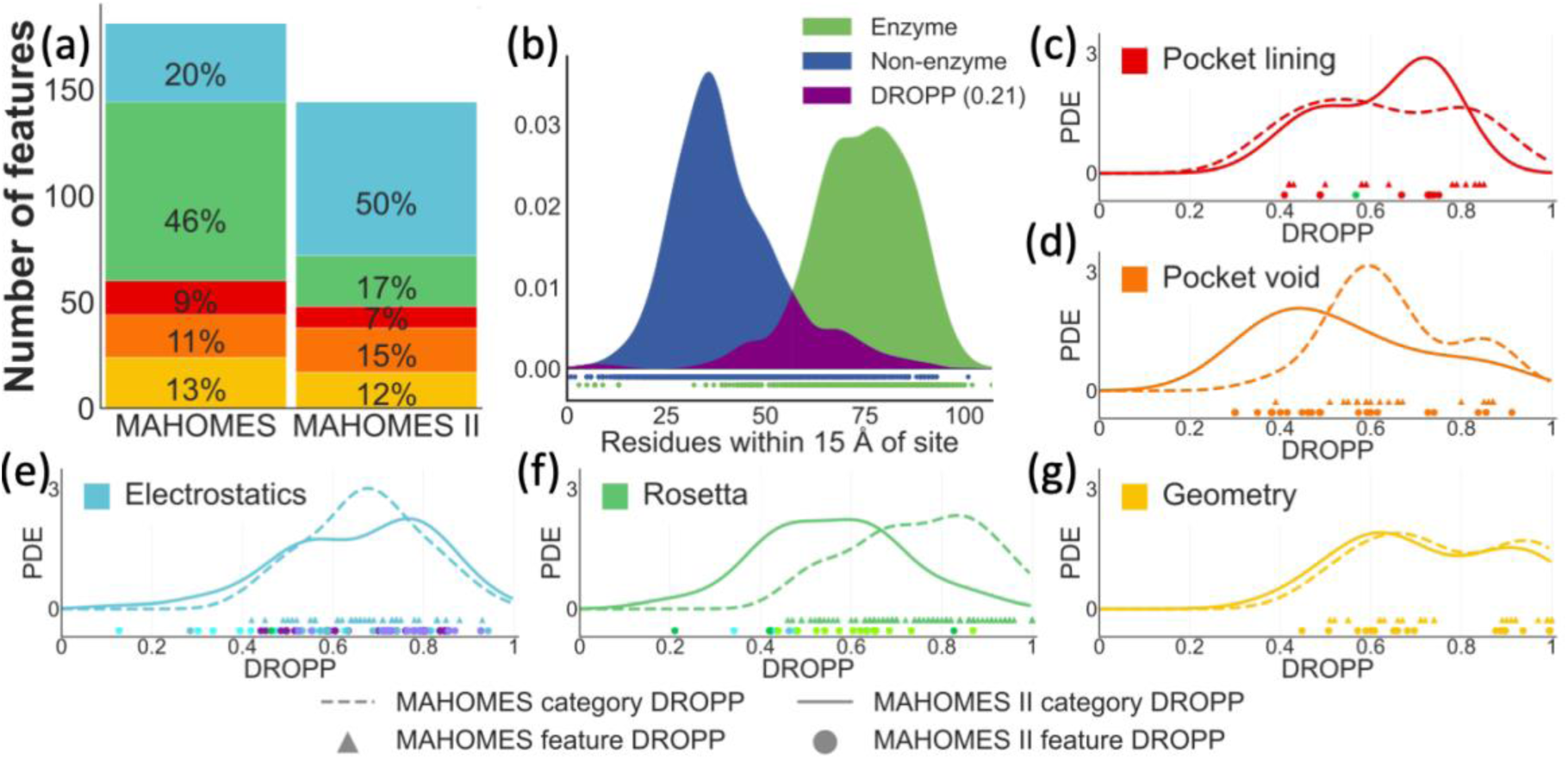
Feature category input space and DROPP. (a) The number of features used by MAHOMES and MAHOMES II for each feature category: blue for electrostatics, green for Rosetta energy terms, red for pocket lining, orange for pocket void, and yellow for coordination geometry. (b) Example of DROPP calculation for the number of residues within 15 Å of the site feature. Kernel density estimators for the feature’s dataset values of enzymes (green) and non-enzymes (blue) are made and the overlapping region (purple) is calculated to give the features DROPP. (c-g) Comparison of MAHOMES and MAHOMES II DROPP probability density estimate (PDE) for each feature category:(c) pocket lining category, (d) pocket void category, (e) electrostatics category, (f) Rosetta energy terms category, and (g) geometry category. Dotted lines and triangles represent MAHOMES features. Solid lines and circles represent features used by MAHOMES II. Circles are colored by feature groups shown in figure 4b.

We use a metric to quantify how much a feature’s values are similar between enzymatic and non-enzymatic sites. This metric, DROPP (<u>d</u>istribution <u>o</u>verlap of a <u>p</u>hysicochemical <u>p</u>roperty) identifies how similar the distribution is between enzymatic and non-enzymatic sites(Figure 1b)(see methods). DROPP was previously found to be lower for features that are more important for predicting enzyme sites.(Feehan, Franklin, and Slusky 2021). Therefore, when trying to improve our features, we used DROPP as indicator of feature improvement (Figure 1 c-g). We made efforts to improve all feature classes except coordination geometry, though some improvements were more successful than others.

#### Electrostatics features expansion

The most important feature used by the original MAHOMES model was an electrostatic feature (Feehan, Franklin, and Slusky 2021), which was the mean second moment of the of the theoretical titration curve’s first derivative for ionizable residues in the second shell (3.5-9Å). We modeled this feature after the THEMATIC calculations, which have been used to identify catalytic residues due to their deviations from Henderson Hasselback titration behavior (Somarowthu, Yang, et al. 2011; Tong et al. 2009; Ko et al. 2005).

To improve our enzyme activity predictions and further our understanding of electrostatic properties responsible for catalytic activity, we expanded our electrostatic features category from 37 features to 152 features (Sup. Figure S1). To further investigate the success of electrostatics in MAHOMES, we added the Z-score calculations which are used by THEMATICs to measure the relative deviation of the theoretical titration curve’s first derivative’s second, third, and fourth moments (Ko et al. 2005). We also added variables output by the generalized Born program we use for generating theoretical titration curves, BLUUES (Fogolari et al. 2012b). Moreover, since catalytic residues often show interesting shifts in pKa (Pérez-Cañadillas et al. 1998; Bate and Warwicker 2004), we added features for the pKa shift from ideal amino acid values, the pKa shift due to desolvation, the pKa shift due to the interaction with other charges in the molecule with all titratable sites in their neutral state, and the pKa shift due to the interaction between titratable sites. Additional added features in the electrostatic category are the generalized Born atomic radii, a solvation exposure parameter, and solvation energies. After removing redundant features (see methods), 72 electrostatic features are used by MAHOMES II (Figure 1a). Six of these new features had lower DROPP than any of the 37 previously used electrostatic features (Figure 1e).

#### Rosetta features reduction

In contrast to the expansion of the electrostatic feature space, we reduced the Rosetta feature space while also improving the features and improving our model reproducibility. We calculated Rosetta features in MAHOMES based on spheres with defined radii from the center of the site. In benchmarking MAHOMES, we found that sub-angstrom differences between relaxed structures of the same site caused large shifts in Rosetta feature values. To prevent sub-angstrom differences from significantly changing calculated feature values for the same site, we switched to a radial basis function (RBF) calculation for the Rosetta energy term features. The RBF calculation uses distance to weight each residue’s influence on the calculated feature, which prevents subtle changes in the structure from having a significant impact on the calculated value. The RBF Rosetta energy terms category decreased DROPP (Figure 1f) and the number of features used by the category (Figure 1a).

#### Pocket void and pocket lining improvements

Our previous method, Rosetta pocket measure (Johnson and Karanicolas 2013), did not detect surface pockets for 645 dataset sites, therefore 19% of the MAHOMES training data was missing values for pocket void and pocket lining features. GHECOM (Kawabata 2019, 2010), a tool that uses mathematical morphology for finding multi-scale pockets on protein surfaces, generated pocket for 99.5% of the dataset sites. To improve the quality of training data, we removed dataset sites that did not have pocket. Additionally, we added various pocket descriptors, including output features from GHECOM which describe the pocket’s shallowness and size rank relative other pockets on the structure. Ultimately, the pocket output by GHECOM lowered the DROPP for features in both the pocket lining and pocket categories (Figure 1c and 1d).

### 2.2 Reduced training data labeling error

Using manual validation, we previously estimated that ∼6% of our non-catalytic sites are mislabeled and that ∼0% of our catalytic sites were mislabeled (Feehan, Franklin, and Slusky 2021). When using cross validation to evaluate newer (intermediate) iterations of MAHOMES, we found seemingly-incorrect predictions were often actually the sites our data generation pipeline mislabeled. We therefore intentionally used ML to hunt for mislabeled sites in our dataset via cross-validation.

Cross-validation is an ML method that leaves out a fraction of the dataset during training so that it can be used to assess the model’s predictive performance. The left-out fraction is iterated over the entire dataset, meaning a model makes predictions for each site in the training dataset. We manually examined non-enzymatic dataset sites that were predicted to be enzymatic during cross validation (see methods for more details). Because manual inspection during work on MAHOMES of 50 random dataset enzyme sites revealed an ∼0% enzyme labeling error (Feehan, Franklin, and Slusky 2021), we did not examine enzymatic sites that were predicted to be non-enzymatic.

We used the available literature (structure publications, RCSB (Burley et al. 2019), and UniProt (UniProt 2019)) to investigate 225 sites that were previously labeled non-enzymatic but were classified during this cross validation as enzymatic. 94 of those sites had definitive literature support of catalytic activity (mislabeled) and 26 PDBs were removed from our set due to inconclusive evidence. Our previous estimate of 6% mislabeled non-catalytic sites implied approximately 158 mislabeled sites in the dataset. Therefore, we estimate that finding 94 mislabeled sites reduces our site mislabeling by 60%.

### 2.3 ML model assessment

We generated 1,792 different ML models (Figure 2) using the following steps: feature standardization, feature selection, and fourteen ML classification algorithms using one of four optimization scoring terms. Since ML algorithms require or are greatly aided by standardization of feature values in order to make comparable scales between the values of different features, we tested four different standardization techniques (see methods). Additionally, large numbers of input features can be detrimental to certain ML algorithms. To decrease the number of features with minimal information loss, we identified four feature subsets each using a different cut off to remove correlated features (see methods). In total we tried six feature sets (four low correlation subsets, all features, and a manually curated set) The six standardized feature sets were then used as inputs to ML classification algorithms which include: linear regression, decision-tree ensemble methods, support vector machines, nearest neighbors, Bayesian classification, and simple neural networks (see methods).

**Figure 2:**
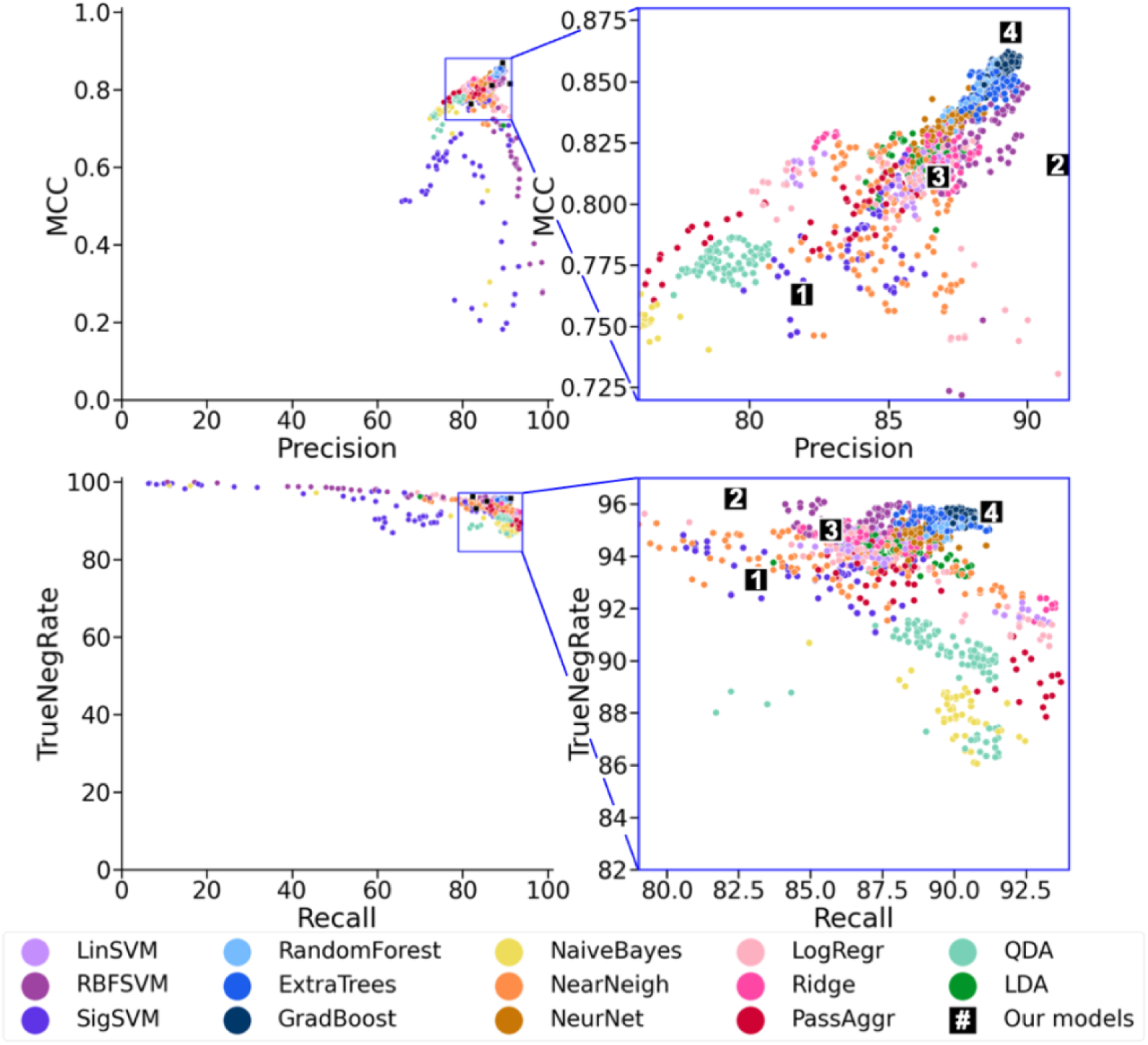
Cross-validation performance by algorithm. Each dot represents one of the 1,792 models assessed in this work. The dots are colored to represent the type of ML algorithm the model uses: support vector machines = purples, decision-tree ensemble methods = blues, linear models = reds, discriminant analysis=greens, naive Bayes = yellow, nearest neighbor = orange, and neural network = brown. Better performing classifiers should have higher precision, Mathews correlation coefficient (MCC), true negative rate (TNR), and recall, meaning better classifiers will be close to the upper right corner. The black boxes with numbers show CV performance of: (1) the previously reported MAHOMES, (2) MAHOMES recalculated with the updated data labels, (3) MAHOMES retrained on updated labels, and (4) MAHOMES II – updated labels, trained on updated labels, and using new features. Right panels are zoomed in views of blue boxes in left panels.

Selecting the best variation of the ML algorithm on the same data used to access a model can inflate performance metrics. To avoid inflated model assessment metrics, we used nested cross validation using an inner loop and an outer loop. During the inner loop, the ML algorithm was fine-tuned for a particular scalar and feature set using one of four different scoring metrics— accuracy, precision, Matthews correlation coefficient (MCC)(Matthews 1975), or a multi-score combination of accuracy, MCC, and Jaccard index(Jaccard 1907). Among our hyperparameter search space, each of the top three ranking ML algorithm variations were used to make models that were accessed using the outer loop. In total, we attempted 4,032 machine learning combinations (14 algorithms x 6 feature sets x 4 standardization techniques x 4 optimization terms x 3 top algorithm variations). Due to convergence during model optimization, this process resulted in 1,792 different ML models.

The vast majority of all attempted ML models in this study outperformed the previous reported MAHOMES cross validation metrics (Figure 2, black box 1) because the training set was substantially corrected for all the new models. In order to make a more fair comparison between MAHOMES and the new models, we re-calculated MAHOMES cross validation performance metrics using the corrected enzyme/non-enzyme labels (Figure 2, black box 2). The number of MAHOMES cross validation false positives dropped from 182 to 90, which increased the precision by nearly 10% (Figure 2, top row) but the rest of the performance metrics remained far below those of our new models.

To assess if the increase in performance was purely due to corrected data labels, we assessed an intermediate model, which retrained MAHOMES using the corrected data but using the old MAHOMES features (Figure 2, black box 3). Despite an increase in recall, the retrained MAHOMES still identified significantly fewer enzyme sites than similar ML models that used the new features (Figure 2, CV blue). Thus, our ML benefitted from the improvement of both the quality of training labels and the improved features.

### 2.4 MAHOMES II performance

To evaluate if these metrics are inflated from overtraining despite cross validation, we also predicted sites in an updated hold-out test set. In addition, we developed a new set derived from the hold-out test set to evaluate the reliability of the models. This set, the T-metal-sites10, includes ten different minimized structures for each site in T-metal-sites. The sub-angstrom variations for each site allowed us to calculate two reproducibility metrics. First, we calculated the divergence frequency (equation 4, methods), which is the percent of test-set sites that received both an enzyme and non-enzyme prediction. Then, we calculated the divergence score (equation 5, methods), a measurement of the severity of divergent predictions. The divergence score ranges from 0 (the site receives the same prediction for every structure) to 1 (the site is predicted enzymatic for half of the structures and non-enzymatic for the other half).

For MAHOMES II, we selected a GradBoost model that used FeatureSet4. We selected that as our final model because of its high cross validation metrics. However, ExtraTrees models, which is the algorithm used by the previous MAHOMES model, had the lowest divergence frequency (Figure S2). So, we further refined hyperparameters, which were too computationally expensive to optimize for GradBoost during our inner cross validation using GridSearch optimization to mimic those favored by the ExtraTrees models (supplemental methods MAHOMES II, fine tuning). The final MAHOMES II model had a cross validation MCC higher than any other ML model (Figure 2 black box 4). Though this could have indicated an overfit model, our final performance evaluation on the hold-out test set T-metal-sites10 (Table 1), which we only used for reproducibility metric calculations during optimization, fell within the projected performance of the CV assessment (Sup. Table S1), thereby supporting the dependability of both the CV assessment and final evaluation of MAHOMES II performance.

**Table 1.** ML model performance evaluations. Predictive performance of MAHOMES, retrained MAHOMES with corrected labels, and MAHOMES II on the holdout test-set, T-metal-sites10, and the quality AlphaFold set, which is the subset of generated sites with confidence scores recommended to characterize binding (pLDDT>90) within 15 Å of the metal. TNR is the true negative rate and MCC is the Mathews correlation coefficient. Descriptions of performance metric calculations in methods. *Evaluation using T-metal-sites, which includes ten incorrectly labeled sites and eight undeterminable sites which were removed from T-metal-sites10.

| ML model | Evaluation | Accuracy | Precision | Recall | TNR | MCC | div. freq. | div. score |
| --- | --- | --- | --- | --- | --- | --- | --- | --- |
| MAHOMES* | T-metal-sites | 94.2 | 92.2 | 90.1 | 96.2 | 0.87 | - | - |
| Recalculated MAHOMES | T-metal-sites10 | 92.6 | 94.2 | 85.3 | 96.9 | 0.84 | 6.6 | 0.53 |
|  | Quality AlphaFold set | 92.7 | 94.1 | 66.7 | 99.0 | 0.75 | - | - |
| Retrained MAHOMES | T-metal-sites10 | 93.4 | 91.9 | 90.0 | 95.4 | 0.86 | 5.9 | 0.55 |
| MAHOMES II | T-metal-sites10 | 94.9 | 94.1 | 92.1 | 96.6 | 0.89 | 5.0 | 0.47 |
|  | Quality AlphaFold set | 97.6 | 95.7 | 91.7 | 99.0 | 0.92 | - | - |

In addition to improved performance, our aim was to lessen the effects of sub-angstrom deviations in input structures. Retraining MAHOMES with corrected labels decreased the divergency frequency but increased the divergence score (Table 1). Updated features in MAHOMES II further decreased the divergence frequency *and* decreased the divergence score, demonstrating that our feature improvements were effective at improving reproducibility.

To test our hypothesis that upgraded features and improved training data can be used to successfully predict enzyme activity for predicted structures, we tested MAHOMES II on a set of AlphaFold generated structures (Tunyasuvunakool et al. 2021; Jumper et al. 2021). However, AlphaFold generated structures do not have ligands such as metals. To create the AlphaFold set of protein structures with metals we queried UniProt (UniProt 2019) for proteins with known metal coordinating residues and no solved crystal structure and filtered for metal ions that could be mapped to AlphaFoldDB structures. Benchmarking our metal method using dataset sites revealed sub-angstrom placement accuracy.

AlphaFold predictions have a confidence metric associated with each residue. The AlphaFold authors recommend using residues with high confidence (pLDDT >90) for characterizing binding sites. Very few sites in our AlphaFold set had high confidence for all residues used to calculate the MAHOMES II features (residues within 15 Å of the metal). So, we made MAHOMES II predictions on the entire non-homologous, metalloprotein AlphaFold set and made multiple performance evaluations by requiring residues within X Å of the site to be high quality (Figure 3, Table 1 and S2). For the entire AlphaFold set, MAHOMES II was able to correctly identify 87% of the enzyme sites (recall) and 90% of the non-enzyme sites (true negative rate)(Sup Table S2). As we removed structures with low quality residue predictions close to the metal, MAHOMES II performance increases up to an accuracy of 97.5% an improvement even over our test-set metrics (Figure 3).

**Figure 3.**
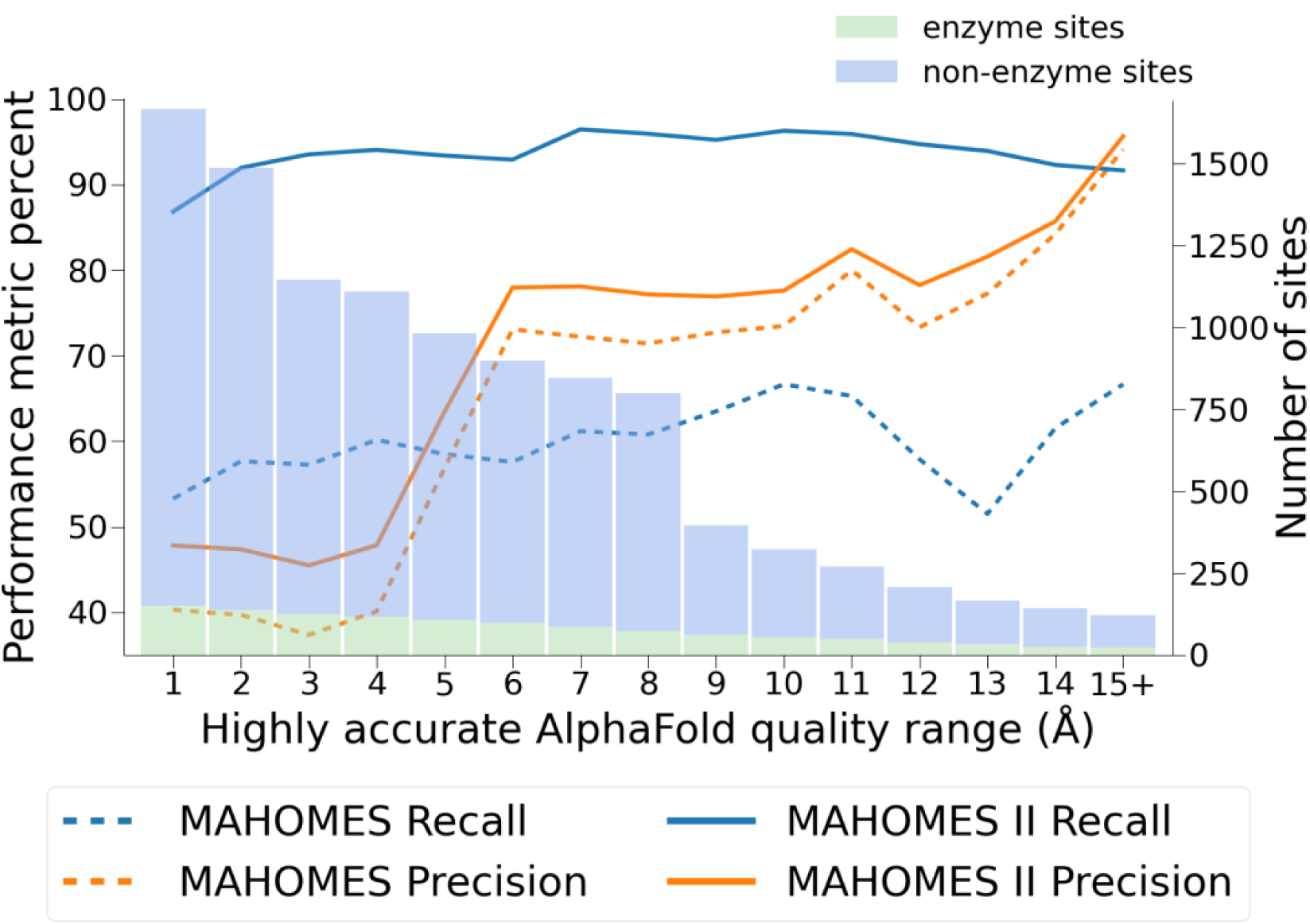
AlphaFold set performance evaluation. The number of enzyme (green bar) and non-enzyme (blue bar) Alphafold set sites containing only highly confident residues within X Å, where X ranges from 1 to 15. The recall (blue lines) and precision (orange lines) of MAHOMES (dotted lines) and MAHOMES II (solid lines) are shown for the Alphafold set sites at each cutoff.

Interestingly, the coordinating residues’ quality was not the most important, as MAHOMES II performance increases the most as the quality range increases from 4Å to 6Å (Figure 3). MAHOMES II enzyme recall (predicting which protein sites are catalytic) was very stable over all confidence regions and vastly outmatched the previous MAHOMES model (Figure 3).

### 2.5 Feature importance

Because MAHOMES II uses a decision-tree based gradient boosting algorithm, we can measure each feature’s importance via the feature contribution to the decrease in impurity on the training data. As previously shown (Feehan, Franklin, and Slusky 2021), features with high importance had low DROPP (overlap between enzyme site and non-enzyme site feature values). The five most important features for MAHOMES II had lower DROPP than any MAHOMES feature (Figure 4). However, the feature with the lowest DROPP, the minimum solvent exposure parameter for outer sphere (3.5 – 9 Å) ionizable residues, was not important to MAHOMES II—it ranked 116^th^ in feature importance for MAHOMES II (Sup Table S3). Hence, though quantitative differences, such as those measured by DROPP, can indicate potentially important features, MAHOMES II is learning more than just these numerical differences in order to successfully differentiate between enzyme and non-enzyme sites.

**Figure 4.**
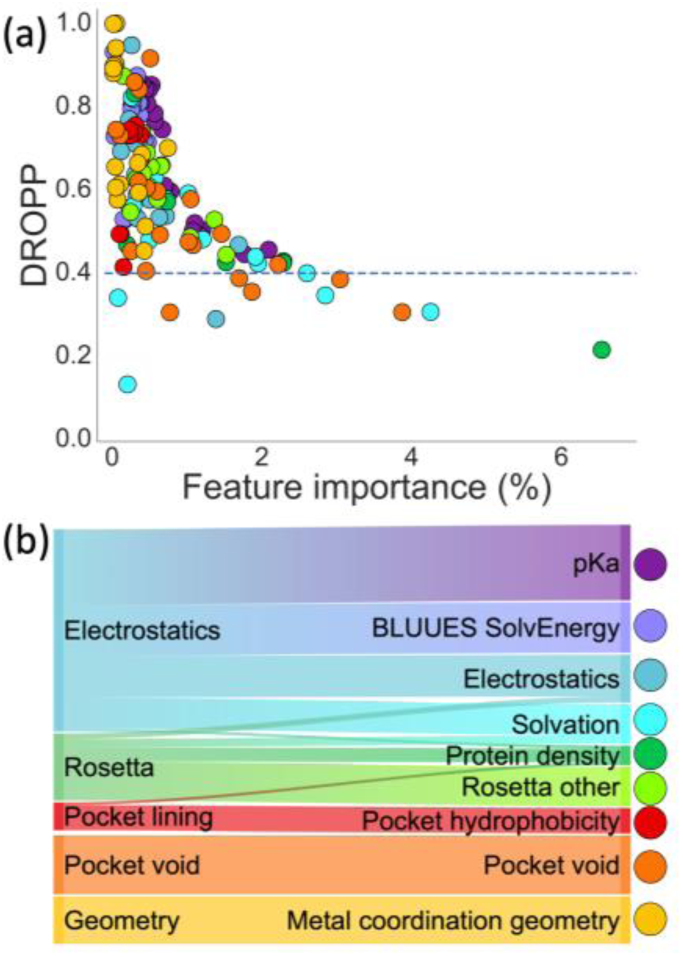
Feature importance and DROPP. (a) Each dot represents MAHOMES II feature and is colored by physicochemical group. The y-axis is the feature’s DROPP, or overlap between values for enzyme and non-enzyme dataset sites. The x-axis is feature importance for MAHOMES II, which is a measurement of the mean decrease of impurity by a feature during training. The blue dotted line represents the lowest feature DROPP from MAHOMES. (b) Sankey diagram of feature distribution between feature categories and feature groups, where width is representative of number of MAHOMES II features.

Since the original feature categories were based on calculation method, we transitioned to feature groups (Fig. 1B) for analyzing which physicochemical properties were the most important for identifying catalytic activity. For example, the Coulombic electrostatic potential RBF feature had been in the Rosetta category but was a better fit for the electrostatic group. Due to differences in feature importance, we split features describing solvation into two groups. The BLUUES SolvEnergy group includes features calculated directly from the BLUUES solvation energy output. We placed other solvation related features in the Solvation group.

The three most important feature groups are local protein density, solvation, and pocket void (Table 2). Despite only making up 29% of MAHOMES II feature space, these groups account for 55% of what the model learned during training. These feature groups also have the lowest average DROPP. Using the DROPP plots for features in these groups, we identify specific subgroups that were fundamental to MAHOMES II distinguishing between enzyme and non-enzyme sites.

**Table 2.** Feature group importance. Each feature group is described by its number of included features (num total), the percent of MAHOMES II feature space accounted for by the group, the total feature importance for all group features, the mean feature importance of features in the group, the rank of the most important feature in the group, and the mean DROPP of features in the group.

| Feature group | Number of features | MAHOMES II feature space | Feature importance |  |  | DROPP mean |
| --- | --- | --- | --- | --- | --- | --- |
|  |  |  | total | mean | Max (rank) |  |
| Local protein density | 7 | 4.9% | 14% | 2.0% | 6.5% (1) | 0.48 |
| solvation | 14 | 9.7% | 18% | 1.3% | 4.3% (2) | 0.46 |
| Pocket void | 21 | 14.6% | 22% | 1.1% | 3.9% (3) | 0.54 |
| pKa | 27 | 18.8% | 19% | 0.7% | 2.1% (10) | 0.69 |
| Rosetta | 14 | 9.7% | 9% | 0.6% | 1.5% (17) | 0.62 |
| Electrostatics | 17 | 11.8% | 9% | 0.5% | 1.7% (16) | 0.63 |
| Pocket hydrophobicity | 9 | 6.2% | 2% | 0.2% | 0.4% (75) | 0.64 |
| BLUUES SolvEnergy | 18 | 12.5% | 4% | 0.2% | 0.5% (59) | 0.75 |
| Metal coordination geometry | 17 | 11.8% | 3% | 0.2% | 0.7% (34) | 0.74 |

The local protein density feature group (seven features) has the highest average feature importance and includes Lennard-Jones energies and the number of residues within a certain distance of the site. This group includes the most important MAHOMES II feature, the number of residues within 15 Å (Figure 1B), which is more important than the 44 least important features combined (Table S3).

The next most important group, the solvation feature group (fourteen features), includes Rosetta solvation features and BLUUES generalized Born features. The second most important MAHOMES II feature is the average BLUUES solvent exposure parameter for second shell (3.5-9Å) ionizable residues. The DROPP plot for this feature shows that most second shell ionizable residues are buried for enzyme sites and relatively exposed for non-enzyme sites (Figure S3b). This group also contains the sixth most important feature, the maximum second shell generalized Born radius, which measures an atom’s shielding from high solvent dielectric (Figure S3f). Enzyme sites also have higher Rosetta solvation features that rank fifth, twelfth, and twenty-first in feature importance (Figure S3e, Table S3), which corresponds with the energetic cost associated with buried charged residues.

The third most important feature group is the pocket void group (twenty-one features). The pocket void group has features that describe the pockets’ location, shape, and size. The third most important MAHOMES II feature describes the slice of the pocket closest to the metal as being larger for enzymes (Figure S3c). The fourth most important feature is the shortest distance between a metal and pocket grid point, which is smaller for enzyme sites (Figure S3d). These features combine to make a subgroup describing site accessible pockets.

## 3. Discussion

Our previous classifier, MAHOMES, outperformed available, alternative methods for classifying enzymes or non-enzymes. MAHOMES II, outperforms its predecessor with increased reliability thanks to both upgraded features and reduced training data error. MAHOMES II’s performance generalize to new, unseen metalloproteins. Moreover, MAHOMES II learned physicochemical properties related to our current understanding of enzyme function.

### 3.1 MAHOMES II learned general enzyme activity

A key question of any classifier is if it has learned beyond its training, i.e. can it predict for examples it has never seen before. For MAHOMES II, training on solved, crystal metalloproteins structures could limit its performance to the 0.056% of proteins with experimentally determined structures (UniProt 2019). Our evaluation using the newly curated AlphaFold set finds that MAHOMES II generalizes to new enzyme reactions and even generalizes to very unrelated proteins.

Alternative tools that can be used to identify enzymatic activity (Zou et al. 2019; Kumar and Skolnick 2012) are less successful than MAHOMES II at predicting if our set of metalloproteins are catalytic (Table S4). Despite using ML, these enzymatic activity classifiers and catalytic residues predictors (Somarowthu and Ondrechen 2012; Song et al. 2018) rely on homology-based features causing their performance to not be transferable to catalysis more generally or be applicable for novel or designed enzymes.

To make our training data different enough from our testing data to facilitate generalizability, our training datasets and test-sets in both in this work and our previous work (Feehan, Franklin, and Slusky 2021) remove redundancy using local similarity. We only kept sites with dissimilar surrounding amino acid identities preventing training and evaluation of repeated sites among homologs and rare cases of similar active sites on different structural folds (Parasuram et al. 2016).

Using the AlphaFold data set, we determined that our model was extremely generalizable and was not implicitly using homology trends. The extensive quantity of AlphaFold structures and experimental data from UniProt for enzyme labeling (instead of homology) allowed us to use a very high E-value of 1, i.e. only proteins with no evolutionary relationship, for creating our AlphaFold set. In comparison, only 17% of our previous test set, T-metal-sites10, sequences have no detectable homology to metalloproteins used for training MAHOMES II (E-value > 1). Furthermore, only seven of the 46 biochemical reactions included in the AlphaFold set are also included in the dataset used to train MAHOMES II. Despite the use of computationally generated structures and strict redundancy removal, MAHOMES II’s 90-97.5% accuracy on the AlphaFold set was similar to its CV and T-metal-sites10 evaluations. Therefore, we believe our assessment of MAHOMES II performance will remain true for any natural metalloprotein structure uploaded by the community on the webserver, even if it is for a novel enzyme reaction. However, due to a lack of available structures, we remain uncertain if MAHOMES II performance transfers to *de novo* metalloproteins.

### 3.2 The less important first shell

Frequently, enzyme bioinformatics focuses on the active site’s first shell, which is the residues interacting directly with substrate(s) or cofactor(s), such as metal ion(s)(Bartlett et al. 2002; Furnham et al. 2016; Ribeiro et al. 2018). The crucial roles played by first shell residues are well supported by conservation and experimental studies (Morley and Kazlauskas 2005; Ribeiro et al. 2020). MAHOMES II has 60 features that describe only first shell properties, covering coordination geometry, inner shell electrostatics ( < 3.5 Å from metal), and pocket lining. Despite making up 42% of MAHOMES II’s feature space, first shell features account for only 18% of feature importance. Since the same metal and coordinating residues are found to participate in enzyme and non-enzyme functions (Lee et al. 2019), it makes sense that first shell features are largely incapable of differentiating enzyme and non-enzyme sites in metalloproteins since in both the first shell coordinates metals. Consequently, despite the well-known critical roles of the first shell, distinction between metallo-enzymes and metallo-proteins is driven by more distant physicochemical properties.

### 3.3 Comparing important MAHOMES II subgroups to current enzyme paradigms

The physicochemical features most important to MAHOMES II success can be considered as three groups/subgroups–1) local protein density, 2) second shell ionizable residue burial, and 3) site accessibility of pockets (in the pocket void feature group) – align with the current paradigm of the enzyme function, which also consists of three features: 1) local environment control of functional sites through control of water access, 2) networks of residue interactions spanning from functional residues, and 3) conformational dynamics (Mazmanian, Sargsyan, and Lim 2020; Agarwal 2019).

#### Control of water access

MAHOMES II captures local environmental control through water access with two of the important MAHOMES II feature subgroups: the local protein density group and site accessibility of pockets feature subgroup. The local protein density features, detect the dense packing of enzyme sites which protects them from high external dielectrics of bulk water, enhancing the local electrostatic effects from hydrogen-bonding and charge-charge interactions. The site accessibility of pockets subgroup identifies close pockets with large openings adjacent to the site that can enable access by individual water molecules, which commonly participate as nucleophiles, to form hydrogen-bonding networks, and to facilitate the release of products. Hence, MAHOMES II can detect the control of water access to enzyme sites by combing local protein density and site accessibility of pockets.

#### Networks of connected interactions

Distal residues that interact with catalytic residues or as part of networks connecting to the catalytic site are essential for fine tuning and optimization of enzyme activity(Dudev et al. 2003; Somarowthu, Brodkin, et al. 2011; Parasuram et al. 2018; Brodkin et al. 2015; Tiwari et al. 2014; Coulther, Ko, and Ondrechen 2021; Coulther et al. 2021; Ngu et al. 2020). The MAHOMES II burial of ionizable residues feature subgroup differentiates enzyme sites based on buried second shell polar and charged residues, which would be a direct result of crucial coupled interactions that enhance enzyme activity. In addition, the MAHOMES II local protein density group uses the density of residues surrounding the active site to provide the most basic description of networks of interactions connected to enzyme sites with the potential to promote activity. The combination of these two subgroups therefore seems to accurately estimate connected networks.

#### Conformational flexibility

Although we did not design any MAHOMES II features to directly describe conformational dynamics, the final aspect of the current enzyme function paradigm, all of the three most important physicochemical subgroups describe properties that affect conformational stability. Local protein density describes tight packing that increases backbone hydrogen-bonding which increases stability and rigidity. Burial of charged residues amongst nonpolar sidechains makes for an energetically unfavorable conformation that will promote destabilization and flexibility. Moreover, interactions between charged sidechains will also increase or decrease stability of various active site conformations depending on the charges. Finally, the site accessibility of pockets enables active site interactions with cofactors, substrates, and solvent that will change the flexibility or rigidity of an active site. Therefore, all three important subgroups contribute to the conformational changes required for enzyme activity, such as the shifting from the ground state to transition state(s).

### 3.4 Machine learning lessons for metalloenzyme design

Considering our training dataset covers all enzyme reaction types(Feehan, Franklin, and Slusky 2021), the physicochemical properties highlighted by MAHOMES II gives us insight for making better metalloenzyme designs. Their feature importance indicates a fundamental blueprint that is harnessed by a range of known catalysis mechanisms performed by nature. To this point, recent work exploring the functional space of non-metallo TIM barrel enzymes has also highlighted the importance of local atomic density(Lipsh-Sokolik et al. 2023). *De novo* enzyme designs on non-enzyme protein backbones could benefit from selecting densely surrounded positions with large pocket openings. Furthermore, lower solvation penalties for buried for ionizable residues might also help design active sites that more closely resemble those in nature. We anticipate that dense protein regions with buried ionizable residues can improve the success rate of designed enzymes and limit additional steps that are currently necessary to reach native enzyme reaction rates, such as directed evolution (Yang, Wu, and Arnold 2019).

## 4. Conclusion

Our ML classifier, MAHOMES II (https://mahomes.ku.edu), uses protein structure-based features describing the local site to distinguish between enzyme and non-enzyme metal ion sites on proteins with 94% precision and 92% recall. We demonstrated that MAHOMES II can make quality predictions for computationally generated structures, which greatly expands its utility when combined with the structure prediction tool AlphaFold. Additionally, the similarity among performance metrics for our cross-validation, holdout test-set, and evolutionarily unrelated AlphaFold set supports that MAHOMES II evaluation is not bias, overfit, or the result of off-target learning. Finally, we were able to identify that MAHOMES II was making successful predictions due to its use of physicochemical features related to densely packed active sites, burial of second shell ionizable residues, and site accessible pockets.

## 5. Methods

### 5.1 Metal ion dataset and T-metal-sites10

The data developed to train and evaluate MAHOMES (Feehan, Franklin, and Slusky 2021) is, to our knowledge, the largest non-redundant dataset of enzymatic and non-enzymatic labeled protein bound metal ions. Briefly, protein structures containing transition metals were filtered to remove poor quality structures and structures dissimilar to metalloenzymes. Metal ion sites bound to multiple chains were removed to avoid labeling partial enzyme active sites during our homology-based enzyme labeling. Metalloenzymes were identified using explicit enzymatic annotations and homology to entries in the Mechanism and Catalytic Site Atlas (M-CSA)(Ribeiro et al. 2018), a database of enzyme active sites. Alignment with M-CSA homolog structures was used to label metal sites on metalloenzymes as enzyme or non-enzyme. Metals on metalloproteins that lacked explicit enzymatic annotations and had no M-CSA homolog were labeled as non-enzyme sites. Finally, sequence and structural homology were used to remove redundancy. Sites on structures deposited in the Protein Data Bank (PDB)(Berman et al. 2000) prior to 2018 were placed in the dataset, which was used for training ML models, ML optimization, and model selection. Sites on structures deposited in the PDB in 2018 or later were placed in a holdout test-set, called T-metal-sites, which was used for final model evaluation.

All the metalloprotein structures in the dataset and T-metal-sites were relaxed using Rosetta (Conway et al. 2014) using a previously provided RosettaScript (Feehan, Franklin, and Slusky 2021). To remove loosely bound metals that are less likely to be physiologically relevant, i.e. crystal artefacts, we removed 729 sites that moved more than 3 Å from the aligned crystal structure. We also removed 179 sites that failed MAHOMES feature calculations since this was commonly due to issues like lack of multiple coordinating atoms. New sites were defined as any metals within 5 Å of each other in a relaxed structure. Since the original dataset defined sites using the crystal structures, the revised set slightly differs in the number of sites. All sites containing a metal atom that was previously a part of an enzyme site were labeled enzyme. Any remaining site with a metal atom that was previously included in a non-enzymatic site was labeled non-enzymatic. All other sites on these structures were discarded. We found that MAHOMES was susceptible to making different predictions for the same site on different relaxed structures. To check model prediction reproducibility, we included the ten relaxed structures with the lowest Rosetta energy units for each metalloprotein in T-metal-sites, making T-metal-sites10. We repeated the labeling procedure for the T-metal-sites10 sites.

We removed sites that were flagged by our automated feature process as problematic or that had extreme outlier feature values (greater than ten standard deviations from the dataset mean). Manual examination of sites with incorrect ML predictions identified a significant number of incorrect non-enzyme labels by our pipeline for sites in both the dataset and T-metal-sites10 (Table S5). Sites found to actually be enzymatic were relabeled and sites with undeterminable enzyme activity were removed. At the end of this work, the MAHOMES II dataset contained 957 enzyme sites and 2,467 non-enzyme sites. The final T-metal-sites10 consisted of 1,895 enzyme entries and 3,277 non- enzyme entries, which were representative of 189 enzyme sites and 328 non-enzyme sites.

### 5.2 Feature engineering

For machine learning input features, we calculated physicochemical properties for five categories – Rosetta energy terms, pocket void, pocket lining, electrostatics, and coordination geometry. A complete feature list with descriptions can be found in Table S6. For exact calculations, please see our available github code(Feehan et al. 2022).

#### 5.2.1 Rosetta energy terms

Rosetta 3.13 was used to score all residues in a metalloprotein structure for all energy terms in the energy function *beta_nov16* with all weights set to one(Alford et al. 2017). For each energy term, *E*, a squared inverse radial basis function (Eq. 1) was used to calculate the energy for a given site, ***s***.

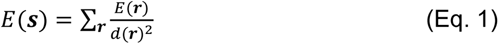

where r is a residue with a distance d(**r**) < 15 Å from the site center. We included the number of residues used for these calculations as a feature. Our Rosetta energy terms category included 25 features in total.

#### 5.2.2 Pocket void terms

We used GHECOM (Kawabata 2019) – a program for detecting multiscale pockets via grid representations and probes – to identify all pockets for a given metalloprotein. Then, for each site on the metalloprotein, we identify all adjacent pockets – pockets with a grid point within 5Å of the site’s center. For sites with multiple adjacent pockets, we select the closest adjacent pocket with a volume greater than 100 Å^3^ as its pocket. In all other cases, the GHECOM pocket with the closest grid point is selected.

We used the selected pocket to calculate the pocket void features previously used by MAHOMES (Feehan, Franklin, and Slusky 2021), which include volume, Euclidean and Manhattan distance between pocket centroid and site center, terms describing the size and shape of three pocket slices at the site center, pocket center, and midpoint of site center and pocket center.

We added pocket void features output by GHECOM for the relative rank among all the metalloprotein’s detected pockets and terms that describe the pockets shallowness and openness of the pocket. Then, we rotated the pocket so that the z-axis runs from the protein centroid to the pockets centroid and calculate its height, max z – min z, and depth, the Euclidean distance between the grid points with the max z and min z. Finally, we calculate the site’s height and depth in the pocket using the Euclidean distance between the site center and the max z grid point or min z grid point respectively.

#### 5.2.3 Pocket lining

The selected GHECOM pocket was used to identify pocket adjacent residues (within 3.0 Å). We identified pocket lining residues as pocket adjacent residues with a sidechain distance of less than 2.2 Å or where the sidechain was closer to the pocket than the backbone. The pocket lining residues were used to calculate the average, minimum, maximum, skew, and standard deviation of the hydrophobicity for both Eisenberg (Eisenberg et al. 1984) and Kyte-Doolittle (Kyte and Doolittle 1982). We also calculated the sum of the pocket lining residues van der Waals sidechain volumes, the volume of the pocket without the lining residues’ sidechains, and the percent of that volume occupied by the sidechains. Finally, the number of pocket lining residues and the number of backbone adjacent residues (pocket adjacent but not considered pocket lining) were included as features.

#### 5.2.4 Electrostatic terms

Our previous electrostatics features were based on the use of theoretical titration curves by THEMATICS (Ondrechen, Clifton, and Ringe 2001; Somarowthu, Yang, et al. 2011; Ko et al. 2005). To calculate these, we used the generalized Borne program, BLUUES (Fogolari et al. 2012a). For the first derivative of the theoretical titration curve, we calculated the second, third, and fourth moment of each ionizable residue using SciPy’s (Virtanen et al. 2020) variation, skew, and kurtosis functions respectively. The mean, standard deviation, maximum, minimum, and range was calculated for two shells. The first, inner shell included ionizable residues within 3.5 Å of a site’s metal atom(s). The second, outer shell included ionizable residues within 9 Å of a site’s metal atom(s), excluding residues that are in the first shell. For each moment calculation, the Z-score was calculated (Eq. 2), where x is the residue’s moment value, μ is the moment’s average for all of the structure’s ionizable residues, and σ is the moment’s standard deviation. We turned this into a site feature by counting the number of residues with a Z-score greater than 1.

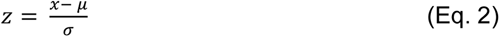

All residues from both shells were used to calculate an environmental feature for each moment using a squared inverse radial basis function (Eq. 1). The number of residues used for the inner shell, outer shell, and environmental feature calculations were also saved to be used as features.

The inner shell, outer shell and environmental features were also calculated for additional BLUUES outputs, which included: generalized Born radius, residuals for the deviation of the theoretical titration curve from a sigmoidal curve, pKa shift from ideal amino acid values, pKa shift due to desolvation, pKa shift due to the interaction with other charges in the molecule with all titratable sites in their neutral state, pKa shift due to the interaction between titratable sites, solvation energies, and solvent exposure parameter. All of the structure’s residues were sorted by BLUUES solvation energy and placed into five bins; destabilizing ranks were assigned from highest to lowest solvation energy and stabilizing ranks were assigned from lowest to highest solvation energy. The inner shell, outer shell and environmental features were calculated for both destabilizing and stabilizing ranks. Overall, there are 152 electrostatic features.

#### 5.2.5 Coordination geometry terms

We used FindGeo (Andreini, Cavallaro, and Lorenzini 2012) to determine the coordination geometries of the site’s metal atom(s). First, we record the total number of ligand N, O, S, and other atoms used as input for FindGeo (within 3.5 Å of any site metal). Then, we record if the metal atom(s) were identified as a regular, distorted, or irregular geometry. If the geometry is regular or distorted, we use the RMSD from the idealized geometry, the number of coordinating atoms for the geometry, and if it is completely or partially filled. To prevent issues with ML algorithms and categorical features, the number of coordinating geometries are one hot key encoded, giving us a total of 20 coordination geometry features.

#### 5.2.6 Feature analysis

DROPP was calculated as previously described for feature similarity (Feehan, Franklin, and Slusky 2021). For discrete features, we used the Jaccard index between the proportions observed in the enzymatic and non-enzymatic sites (Jaccard 1907). For continuous features, we calculated the overlap of the kernel density estimators for the values of the enzymatic and non-enzymatic sites. To prevent extreme outliers from having large influence on DROPP, values greater than ten standard deviations the mean were ignored. Only dataset sites were used for calculating DROPP. The code for the DROPP calculation can be found in the MAHOMES II repository file FeatureCalculations/CalcFeatureDROPP.py (Feehan et al. 2022).

Any feature that had the same value calculated for the entire dataset was discarded, leaving a total of 250 features. To decrease feature space with minimal information loss, additional subsets of features were identified for ML input using maximum correlation cutoffs between features in the same category (Sup. Figure S1). For highly correlated features, the feature with the higher DROPP was removed. FeatureSet2, FeatureSet3, and FeatureSet5 used correlation cutoffs of 0.99, 0.9, and 0.75 respectively. Due to the dramatic increase of electrostatic features, FeatureSet4 used a correlation cutoff of 0.75 for electrostatic features and 0.9 for other categories. Finally, we manually selected features for FeatureSet6.

#### 5.2.7 Preparation for ML

ML algorithms require or are greatly aided by standardization of feature values to take them close to zero and make comparable scales between the values of different features. We selected four different standardization techniques available in scikit-learn (Pedregosa et al. 2011) to use during model optimization and selection. The StandardScaler removes the mean and divides by the features standard deviation. The RobustScaler removes the median and divides by range between the 20th and 80th quantile to mitigate the effect of extreme outliers. We also examined uniform and gaussian QuantileTransformers which use non-linear transformations to map feature values to uniform or gaussian distributions respectively. All scalars include an imputer to fill missing feature values with the dataset sites’ average feature values. MAHOMES II used the uniform QuantileTransformer.

### 5.3 Machine Learning

#### 5.3.1 Classification performance metric calculations

To calculate predictive performance metrics, a prediction is counted as a true positive (TP) if it is an enzyme prediction for an enzyme labeled site. It is considered a false positive (FP) if it is an enzyme prediction for a non-enzyme labeled site. A true negative (TN) is a non-enzyme prediction for a non-enzyme labeled site. Finally, a false negative (FN) is a non-enzyme prediction for an enzyme labeled site. The TP, FP, TN, and FN counts are then used to calculate a model’s accuracy, precision, recall, true negative rate (TNR), and Matthews correlation coefficient (MCC)(Matthews 1975).

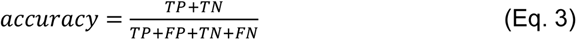

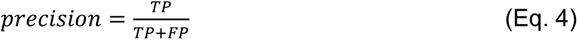

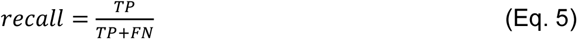

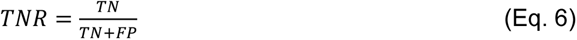

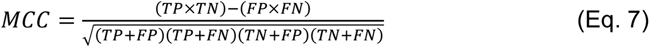

#### 5.3.2 Optimization of ML model(s)

Since the best performing hyperparameters for ML classification algorithm can change depending on input feature subset and standardization technique, a nested cross validation (CV) strategy was used to find the optimal ML models. The outer CV used stratified k-fold and the inner loop used stratified shuffle split. During the inner loop, different hyperparameters sets were attempted and scored using one of four terms - accuracy, precision, MCC, or a multi-score combination of accuracy, MCC and Jaccard index. The hyperparameter sets were ranked according to the average of the scoring metric. Our multi-score optimization ranked each scoring metric and then averaged the rankings. Due to the potential for ties, the top three were selected as the optimized ML models. Depending on the convergence of the different scoring terms, ML algorithm, feature subset, and standardization could have between three and twelve optimized ML models. To reliably compare optimized ML models, we used the average performance metrics during stratified k-fold CV with ten repetitions during each iteration using different random state hyperparameters for the classifier when applicable.

#### 5.3.3 Evaluating a model’s reproducibility

For considering a model’s reproducibility, different minimized structure inputs in T-metal-sites10 were used to make a set of predictions, **p**, for the same site, s. The site’s divergence, d(*s*), was calculated using Equation 8, where *p_i_* is either 1 (enzyme prediction) or −1 (non-enzyme prediction) and *n* is the number of minimized input versions for s.

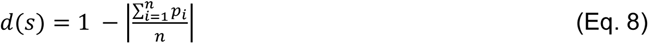

Therefore, d(s) ranges from 0 to 1, where 0 is a site with the same prediction for all minimized inputs and 1 is a site with five enzyme predictions and five non-enzyme predictions. Using the set of all sites in T-metal-sites10, **T**, and the subset of divergent sites, *D* = {*s* | *s* ∈ ***T*** and *d*(*s*) > 0}, we calculated our reliability metrics. The divergence frequency is the percent of sites in T-metal-sites10 that were divergent (Eq. 9).

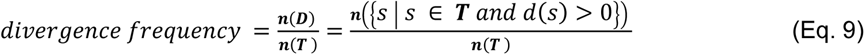

The divergence score is the average site divergence of the divergent sites (Eq. 10).

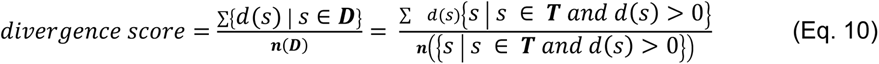

Since only divergent sites are considered, the lowest divergence score is 0.2.

#### 5.3.4 ML guided dataset manual annotation

We optimized decision tree-based algorithms (ExtraTrees, GradBoost, and RandomForest) using FeatureSet4 and selected an ExtraTrees model with high MCC and high recall. We manually checked non-enzyme sites that were predicted to be enzyme sites during the k-fold CV using available structure publications, RCSB(Burley et al. 2019), and UniProt(UniProt 2019). We changed the labels for 73 sites that were actually enzyme sites and removed all sites from 16 metalloproteins that were undeterminable, for reasons such as lack of publication and enzymatic homologs. We repeated the process without ExtraTrees models to minimize redundancy of mispredicted sites. We fixed 19 additional site labels found during the second iteration, removed all sites from eight undeterminable metalloproteins. We also removed two sites that were located on the edge of active sites, meaning they could be correctly labeled as both enzymatic and non-enzymatic.

#### 5.3.5 Recalculated MAHOMES and Retrained MAHOMES

Since we were able to identify a significant amount of labeling error in the data used for previously reported MAHOMES performance evaluations, we made updated performance evaluations to enable more fair comparisons between MAHOMES and MAHOMES II. We recalculated the k-fold performance for the MAHOMES predictions using the corrected dataset labels for what was and wasn’t an enzyme. Since we made new relaxed structures for T-metal-site10 (the MAHOMES II test set), new MAHOMES predictions were made for T-metal-sites10. Both variations of the test-set received nearly the same predictions, but the performance evaluation changes significantly due to reduced labeling error in T-metal-sites10.

The recalculated MAHOMES performance evaluations were still from a model that was trained using dataset sites which have since been identified as mislabeled. So, we made a retrained MAHOMES model, which differs from recalculated MAHOMES in two ways. The first difference is that the fixed dataset with updated labels and removed undeterminable sites were used during training. The number of enzyme sites increases by 10% when the dataset labels are fixed, which prevents under-sampling at a ratio of 3 non-enzyme:1 enzyme site during training. So, the retrained MAHOMES model under-samples by randomly removing 10% of the enzyme sites, followed by random removal of non-enzyme sites until the ratio of training data is 3 non-enzyme:1 enzyme sites. Otherwise, retrained MAHOMES model uses the same methods as the recalculated MAHOMES model, including calculated feature values, algorithm, and optimized hyperparameter set.

#### 5.3.6 Model selection and performance evaluation

Despite the favorable performance of decision tree-based classifiers during work on MAHOMES(Feehan, Franklin, and Slusky 2021), we tested fourteen ML classification algorithms from Scikit-learn(Pedregosa et al. 2011) for MAHOMES II, which include: linear regression, decision-tree ensemble methods, support vector machines, nearest neighbors, Bayesian classification, and simple neural networks. We decided to attempt these various algorithms because decision tree ensemble-based classifiers are known to be robust against mislabeled data, large feature spaces, and outlier feature values. So, our upgraded features and reduced training label error does not affect decision tree ensemble-based algorithms as much as it affects alternative ML classification algorithms.

In total, we assessed 4,032 machine learning combinations (14 algorithms x 6 feature sets x 4 standardization techniques x 4 optimization terms x 3 top hyperparameter sets). However, we only ended up with 1,792 unique ML models due to convergence during model optimization. The specific code used for the algorithms, standardization techniques, and hyperparameters can be found in the MAHOMES II repository file MachineLearning/GeneralML.py (Feehan et al. 2022).

We selected a model that used a gradient boosting classifier with FeatureSet4 and a uniform QuantileTransformer because it had the highest recall for models with greater than 0.845 MCC and 88.5% precision. For MAHOMES II, we further refined this model to improve both its cross validation MCC, divergence frequency and divergence score by adjusting hyper-parameters that were too computationally expensive to optimize during our inner cross validation using GridSearch optimization (Sup. methods).

#### 5.3.7 Feature importance

For algorithms using decision-tree classifiers, sci-kit learn has a built in feature importance output that measures the mean decrease in impurity that a feature was responsible for during training. The MAHOMES II feature importance is the average of feature importance output of the models trained during k-fold cross validation.

#### 5.4 AlphaFold set

#### 5.4.1 AlphaFold set generation

To make the AlphaFold set, we queried UniProt(UniProt 2019) for reviewed entries with no solved crystal structure, an AlphaFold model (as of February 15, 2022), and metal binding data. Entries with no EC number and no catalytic activity annotation of any kind were labeled as non-enzyme. Entries with annotated with experimental catalytic activity were labeled as enzyme. Remaining unlabeled entries were removed.

Since each entry is a protein sequence at this point, we chose to remove homology with training and evaluating data next. We used PHMMER (Eddy 2011) to search each protein sequence against all sequences in the PDB as of May 21, 2020. Entries with detected homology, using an E-value < 1, to any protein sequence in the dataset or test-set were removed.

To go from sequence to the site level data, we retrieved all available metal binding data from UniProt for each of the remaining entries. We removed data for metals other than Copper, Iron, Magnesium, Manganese, Zinc, and Nickel. To ensure that the labels were accurate at the site level, we removed enzyme labeled entries that did not include ‘catalytic’ annotations for the metal binding site. Due to automatic metal site annotations or lack of EC coverage, non-enzyme labeled entries with ‘catalytic’ metal binding annotations also had to be removed. Entries with only one or two listed metal binding residue(s) were removed. We did not relax or perform any additional structure minimization. The resulting AlphaFold set contains 1740 computationally generated structures with 1583 non-enzyme sites and 157 enzyme sites.

We placed metals in sites using the average coordinates of the atoms binding to the metals. For hydrophilic amino acids, we used the coordinate of the sidechain atom capable of binding a metal ion (N, O, or S). For amino acids with multiple sidechain atoms capable of binding metal residues (GLU, ASP, GLN, ASN), we used the average of these atomic coordinates. For GLY, we used the average coordinate of the backbone N and O. Some entries also listed other non-polar amino acids as coordinating residues. We found the average coordinate of sidechain carbons worked best for placing these metals without any steric clash. Since some entries with multiple metal binding sites did not differentiate different bound metal sites, we removed any entries if the coordinating residues where more than 12 Å from each other. The resulting 1,675 metalloprotein structures and enzyme/non-enzyme labels are available on Zenodo (Feehan 2023).

#### 5.4.2 AlphaFold set metal ion placement accuracy

We evaluated the accuracy of the AlphaFold set metal placement by adding a metal to the sites in our dataset and test-set using UniProt data and comparing it to the metal location in the relaxed crystal structures.

To find appropriate crystalographically-resolved sites to compare with the AlphaFold set metal placement, we retrieved available binding site data for 2,608 of the 2,643 UniProt entries in our dataset and test-set. However, only 1,207 entries included data for relevant metal binding sites -- CHEBI ids: 29105 (Zn2+), 29033 (Fe2+), 29034 (Fe3+), 29035 (Mn2+), 29036 (Cu2+), 49552 (Cu+), 18420 (Mg2+), or 49786 (Ni2+). To create the dataset for benchmarking, we removed UniProt entries and PDBs with multiple sites. Also, entries with different metals in the PDB and UniProt binding data were removed. To accurately depict the placement of sites in our AlphaFold set, we removed sites with fewer than three coordinating residues. Finally, coordinating residues had to be among the previously described amino acid types, resolved in the PDB, and indexed with the same numbers in UniProt and the PDB. These filtering steps resulted in a total of 103 sites remaining for benchmarking.

The final benchmark set consists of 103 successfully placed sites. The average distance between the placed metal and the metal in the relaxed crystal structure was 0.87 Å (Fig. S4). For comparison, the same 103 sites moved an average of 0.54 Å during minimization of the crystal structure with Rosetta relax. Moreover, 56% of sites were placed within 1 Å, 96% were placed within 2 Å, and only one was more than 3 Å from its respective experimentally resolved location in the PDB structure.

The UniProt metal binding annotations were converted to binding site annotations (Coudert et al. 2023).This data conversion occurred after we created the AlphaFold set but before we benchmarked our metal binding site placement method. This conversion therefore required different data retrieval scripts for the benchmarking set than for the AlphaFold set.

### 5.5 webserver

The MAHOMES Web Server was implemented in Python 3 on the back end using the Flask framework with Jinja for templates in creating the HTML client-side interfaces. When a PDB file and email are submitted to MAHOMES, metadata about the job is stored in a JSON file. The information necessary to schedule the job for processing is placed into an SQLite3 database.

The job execution program, which is written in Python 3, monitors the SQLite database for new user submissions, and then handles executing the job, monitoring the job execution, and then sending an email to the user with a link to the results page.

The jobs and the web application are run on the Slusky Lab web server, which is a virtual machine running in the University of Kansas’s enterprise data center. Running this service as a virtual machine has allowed us to scale up the hardware backing the instance as we have needed additional resources while working to control the long-term costs associated with running the MAHOMES service.

## 6. Supplementary material description

*Figure S1.* Comparison of feature sets.

*Figure S2.* ML models reproducibility.

*Figure S3.* Top 6 feature DROPP plots

*Figure S4.* Metal binding site placement accuracy

*Table S1.* Performance evaluations.

*Table S2.* AlphaFold set evaluations.

*Table S3.* MAHOMES II feature importance

*Table S4.* Comparison of MAHOMES II performance to similar tools that make enzymatic and non-enzymatic predictions

*Table S5.* Manually annotated sites

*Table S6.* Feature details

## Abbreviations footnote

ML: machine learning
RBF: Radial Basis Function
CV: cross validation
MAHOMES: metal activity heuristic of metalloprotein and enzyme sites
DROPP: distribution overlap of a physicochemical property
PDB: Protein Data Bank
PDE: probability density estimate
pLDDT: predicted Local Distance Difference Test
MCC: Mathews correlation coefficient
TNR: true negative rate
TN: true negative
TP: true positive
FN: false negative
FP: false positive

## 7. Acknowledgements

We gratefully acknowledge helpful feedback from members of the Slusky lab, especially Daniel Montezano and Samuel Lim. We also gratefully acknowledge NIGMS awards DP2GM128201 and P20GM103418 to JSGS, NSF award 2226804 to JSGS, and funding from the Scandinavian American Foundation to JSGS.

## 8. Data and code availability

The data and code used to train and evaluate MAHOMES II can be found at https://github.com/SluskyLab/MAHOMES-II (Feehan et al. 2022). The AlphaFold set metalloprotein structures and enzyme/non-enzyme labels can be downloaded from https://doi.org/10.5281/zenodo.7703098 (Feehan 2023).

## 8. Conflicts of interests

The authors declare no conflicts of interests.

## Notes

https://mahomes.ku.edu

https://github.com/SluskyLab/MAHOMES-II

https://doi.org/10.5281/zenodo.7703098

### Competing Interest Statement

The authors have declared no competing interest.

